# Behavioral traits that define social dominance are the same that reduce social influence in a consensus task

**DOI:** 10.1101/845628

**Authors:** Mariana Rodriguez-Santiago, Paul Nührenberg, James Derry, Oliver Deussen, Linda K Garrison, Sylvia F Garza, Fritz Francisco, Hans A Hofmann, Alex Jordan

**Author notes:** Denotes co-first authorship (Mariana Rodriguez-Santiago, Paul Nührenberg). Author Contributions AJ, JD, and HAH designed experiments; AJ and JD wrote code for automated data acquisition; AJ, MRS, LKG and SFG performed experiments; PN performed computer vision tracking; AJ and SFG performed image-based tracking; PN, AJ, LKG, FF, and MRS performed statistical analysis; AJ, PN, LKG, and MRS wrote the manuscript; AJ, HAH, and OD revised manuscript.

## Abstract

In many species, cultures, and contexts, social dominance reflects the ability to exert influence over others, and the question of what makes an effective leader is pertinent to a range of disciplines and contexts. While dominant individuals are often assumed to be most influential, the behavioral traits that make them dominant may also make them socially aversive and thereby reduce their influence. Here we examine the influence of dominant and subordinate males on group behavior in different social contexts using the cichlid fish *Astatotilapia burtoni*. We find that phenotypically dominant males display behavioral traits that typify leadership across taxonomic systems – aggressive, social centrality, and movement leadership, while subordinate males are passive, socially peripheral, and have little influence over movement. However, in a more complex group-consensus task involving visual cue associations, subordinate males become the most effective agents of social change. We find that dominant males are spatially distant and have lower signal-to-noise ratios of informative behavior in the association task, potentially interfering with their ability to generate group-consensus. In contrast, subordinate males are physically close to other group members, have high signal-to-noise behaviors in the association task, and visual connectivity to other group members equal to that of dominant males. The attributes that define effective social influence are therefore highly context-specific, with socially and phenotypically dominant males being influential in routine but not complex social scenarios. These results demonstrate that behavioral traits that are typical of socially dominant individuals may actually reduce their social influence in other contexts.

**Significance Statement:** The attributes that allow individuals to attain positions of social power and dominance are common across many vertebrate social systems – aggression, intimidation, coercion. Yet these traits are socially aversive, and can make dominant individuals poor agents of social change. In a vertebrate system (social cichlid fish) we show that dominant males are aggressive, socially central, and lead group movement. Yet dominant males are poor effectors of consensus in an more sophisticated association task compared with passive, socially-peripheral subordinate males. The most effective agents of social influence possess behavioral traits opposite of those typically found in position of social dominance, suggesting the behavioral processes that generate social dominance may simultaneously place the most ineffective leaders in positions of power.

## Introduction

In human and many non-human animal groups, hierarchy is considered a defining feature of social interactions (1–4). Within groups, hierarchical differences can influence access to resources, mating opportunities, and patterns of conflict (5, 6). Dominance is also frequently associated with leadership in either movement or opinion (7–9). Dominance structures can modify influence in groups, specifically through differences in attention to dominant and subordinate individuals (10). The behavioral attributes that define social dominance may also directly increase social influence, for example confidence and assertiveness in humans (11), so much so that *social dominance* is often considered to be equivalent to *social influence* (for discussion, see 12). Yet dominance may also be associated with aversive behavioral traits, meaning that individuals able to rise to socially dominant positions are the most damaging for group performance. For example, aggression and competitiveness are traits commonly associated with social dominance, yet can potentially lead to the phenomenon of ‘toxic leadership’ in organizations (13, 14).

Dissociating the factors that interact to mediate dominance and influence is therefore crucial for understanding how social hierarchy affects group performance and function, but manipulative experiments in humans can be ethically and logistically challenging. In contrast, social animals can provide excellent models for examining dominance and influence since social dominance is correlated with or a direct result of, higher aggression (15–19). However, it can be challenging to dissociate the effects of dominance status itself from the behavior of the dominant individual. Group movement may be disproportionally influenced by dominant individuals, potentially due to affiliative social bonds (e.g. 9), but influence over movement can similarly be achieved through repulsive interactions (i.e. by pushing or driving others away), which may be predicted when interacting with aggressive dominant individuals. Thus, the link between the influence of socially dominant individuals and the behaviors of those individuals may be context-specific. While dominance and influence may be associated in competitive scenarios, the same aggressive traits may reduce social influence and be detrimental to group function and cohesion in others contexts (20, 21). The difficulty in separating behavioral elements limits our ability to understand the development, evolution, and expression of behavioral traits and their interactions in social contexts, as well as the adaptive significance of dominance (22, 23) and social influence (6).

Here we examine how consensus responses in a socially-facilitated group association task are influenced by social hierarchy. We explore how the behavioral traits that define social dominance in the male cichlid fish *Astatotilapia burtoni* (24) interact with their influence in different contexts. Dominant males of this species have clear phenotypic signatures of social status; they are territorial, brightly colored, and highly aggressive, while subordinate males are non-territorial, non-aggressive, and cryptically colored. Additionally, the presence of these distinct male types can differently influence the behavior of other group members (25). Depending on social context, males can switch between these phenotypes in as little as 20 minutes when the opportunity for social ascension arises (26, 27). Dominant males also have higher degree centrality than subordinates in behavioral social networks (28), frequently exchanging aggressive displays with other males, attacking and chasing members, and courting females. Using this species, we allow dominant and subordinate males to learn a group association task, then place these individuals into new groups of naïve individuals. We ask how the social status of the informed individual affects the time taken for the naïve group to reach consensus and move as a cohesive group to the correct conditioned stimulus. We hypothesize that socially dominant informants will have stronger influence on group level behavior if these individuals occupy central social network positions and information flows along these network edges. Alternatively, the behavioral traits of dominant males may prove an aversive source of social information and make those males ineffective agents of social influence.

## Results

### Groups with subordinate informants reached consensus faster than those with dominant informants

We examined whether the social influence displayed by dominant males in routine conditions extended to a more complex task. We trained dominant and subordinate males on a basic association task and then placed these informed males into groups of individuals naïve to the association task. We measured the speed with which groups containing informed dominant or subordinates reached consensus in moving to the correct unconditioned stimulus, comparing this to the baseline performance of completely naïve groups (Figure 1a). We found that groups containing dominant male informants did not reach consensus faster than naïve groups (log-rank tests, *dominant informant vs naïve*, Χ^2^(log-rank) = 1.13, *p* = .288; Figure 1b), whereas groups containing a subordinate male informant achieved consensus significantly faster than both naïve and dominant male informant groups (*subordinate informant vs naïve*, Χ^2^(log-rank) = 13.99, *p* < .001; *subordinate informant vs dominant informant*, Χ^2^(log-rank) = 5.94, *p* = .015; Figure 1b).

**Figure 1.**
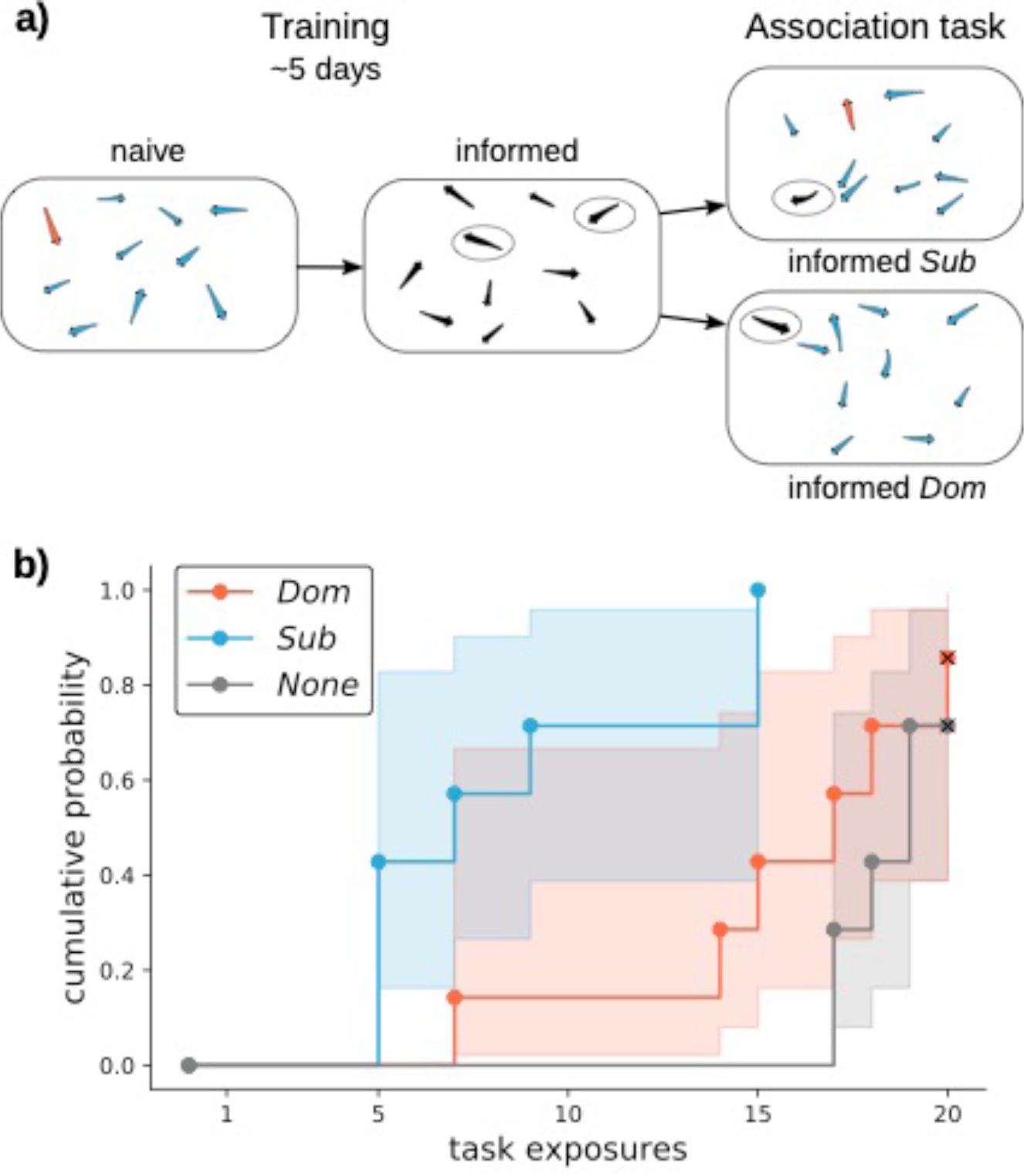
**a)** Experimental protocol of group-consensus association task. Groups of eight individuals (4 males, 4 females, dominant male indicated in this case by orange coloration) that are naïve to the association task are placed into the arena. Over the course of 20 exposures to the association task (4 exposures per day), these naïve individuals become informed about the correct light stimulus (indicated by stippling). After this period, two informed individuals (in one treatment a dominant male, in the other a subordinate male) are moved into demographically identical new groups of naïve individuals. We then measure the speed at which the seven of the eight individuals move towards the correct unconditioned stimulus for two subsequent trials (“group-consensus”). **b)** The cumulative probability (i.e. the inverse Kaplan-Meier probability) of group-consensus in over the course of 20 exposures. Groups that did not complete the task were right-censored in the analysis (indicated by “x”), shaded areas represent 95% confidence intervals. Groups with a *subordinate* (*Sub*) male informant had a higher probability of response than those with a *dominant* (*Dom*) male informant or those without an informant (*None*; log-rank test, *p* < .015).

### Phenotypically dominant males are more central in social networks and lead group movements

We first examined baseline social behavior by placing groups of fish in large holding aquaria and using computer-vision based tracking of interactions to generate social network and behavioral data. Comparing the behavioral interaction network positions of dominant and subordinate males in routine social contexts, we found that dominant males occupy more central social network positions (Figure 2a; Bonferroni corrected Wilcoxon signed-rank tests (α-level = .01); *dominant vs subordinate centrality, Z* = 3.86, p < .01; Figure 2a,b). We then tested whether leadership of routine group movements differed among dominant and subordinate males by analyzing the onset and duration of rapid movement events that dominant and subordinate males either initiated or responded to (example in Figure 2d). We found that dominant males were more effective initiators of group movements than subordinate males in these routine conditions (*dominant vs subordinate motion delay, Z* = -3.66, p < .001; Figure 2c).

**Figure 2.**
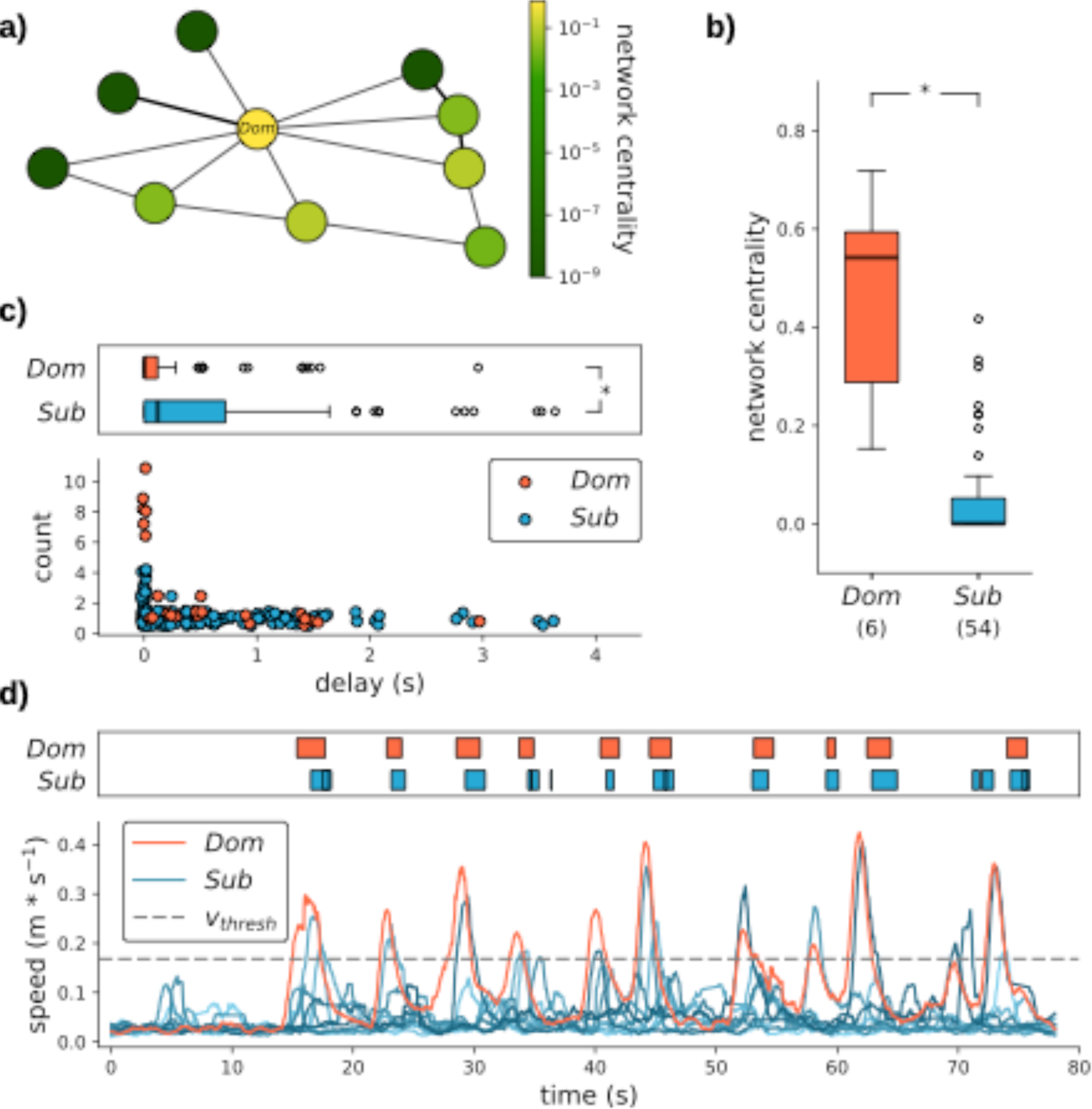
Interaction networks and behavioral analysis from trajectory data. **a)** Example of a social network graph created from directed, pairwise initiator-responder counts (graph directedness omitted for visualization). Node color denotes network (betweenness) centrality of *dominant* (*Dom*) and *subordinate* (*Sub*) group members. **b)** Aggregation of all individuals from the standard condition recordings showing the effect of social status on network centrality (Wilcoxon signed-rank test, *p* < .01). **c)** Onset of speed events across all dominant and subordinate male event initiations (delay = 0) and responses (delay > 0), with upper panel demonstrating a temporal offset in which dominant males more often initiate rapid movements (Wilcoxon signed-rank test, *p* < .001). **d)** Representative examples of speed traces of dominant and subordinate males. Upper panel represents onset and duration of speed events that exceed a 95% threshold of all speeds (‘*v*_thresh_’).

### Dominant males had greater spatial separation and lower behavioral signal-to-noise ratio

We then explored behavioral and social attributes that may have caused this difference in influence across the two social contexts (routine social interactions versus group association task). We first compared the spatial and visual connectivity to other group members during routine social interactions. We found that dominant males were more spatially distant from other group members than were subordinate males (Bonferroni corrected Wilcoxon signed-rank tests (α-level = .01); *dominant vs subordinate mean pairwise distance*, Z = 3.51, *p* < .01; Figure 3b). However, there was no difference in the visual connectivity to other group members between dominants and subordinates (*dominant vs subordinate mean angular area subtended on retina*, Z = -1.44, *p* = .149; Figure 3c).

**Figure 3.**
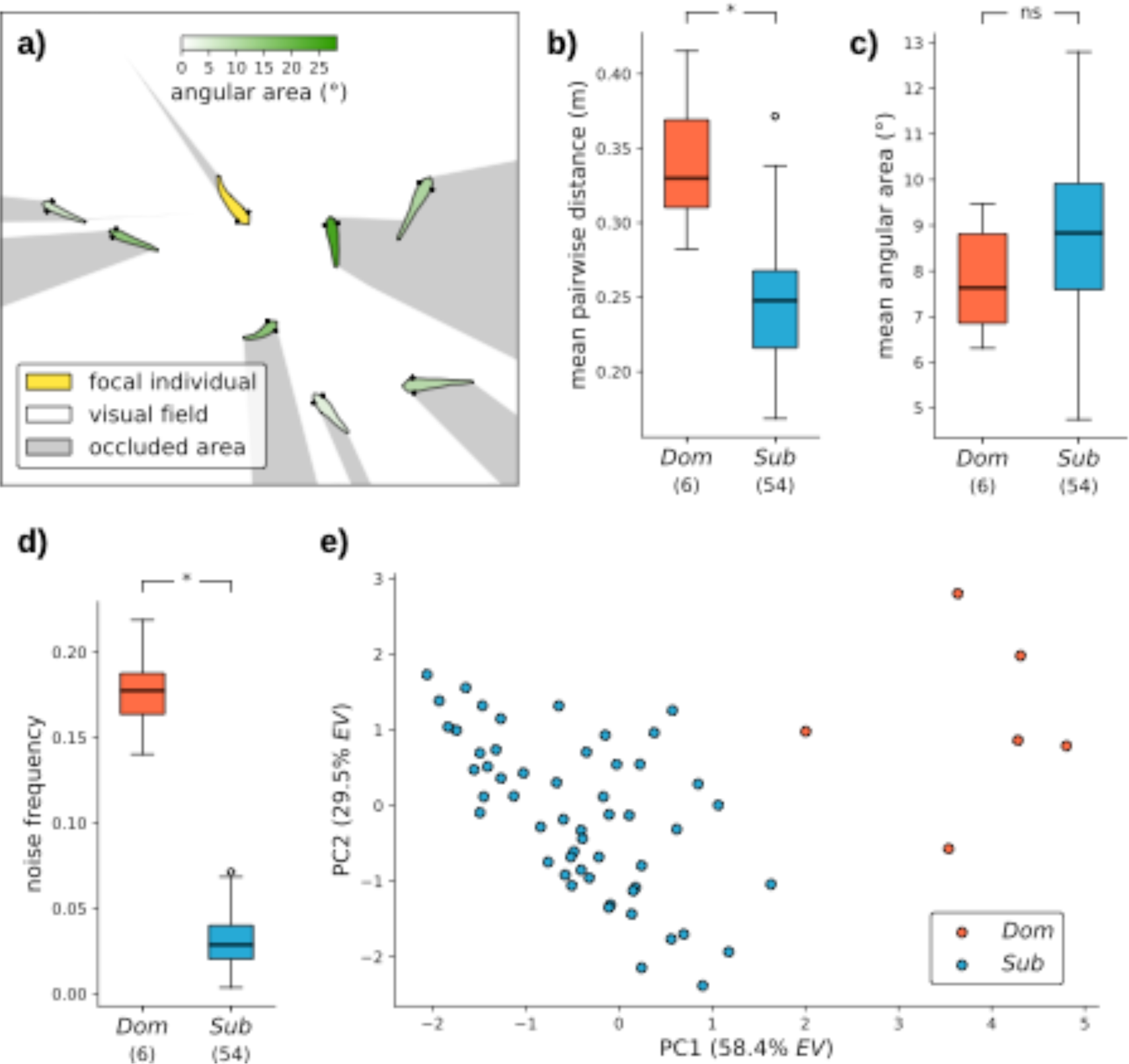
Effects of social status on behavioral parameters. **a)** Schematic of visual field computation. **b)** and **c)** Aggregated data from all standard condition recordings, comparing mean pairwise distance (‘association’ connectivity) and mean angular area (visual connectivity) between *dominant* (*Dom*) and *subordinate* (*Sub*) males with other group members (Wilcoxon signed-rank tests, p < 0.01 and p = 0.149). **d)** Hypothetical signal-to-noise ratio in social learning context (rapid, directed swimming) compared by social status (p < 0.001). **e)** Two-dimensional PCA on all metrics derived from trajectory and network analyses, comparing dominant and subordinate males in this social parameter space. The first two dimensions explain 58.4% and 29.5% of the variance (*EV*).

We then compared the motion signatures of males during routine movement versus during the association task. We found that dominant males were significantly more likely than subordinate males to swim as fast or faster than the average speed of informed individuals moving towards the unconditioned stimulus during the association task, allowing us to generate a predicted signal-to-noise ratio of informative social ‘signal’ (rapid, directed swimming towards correct unconditioned stimulus) versus uninformative social ‘noise’ (rapid, directed swimming during chasing of conspecifics). In this analysis, dominant males had significantly worse signal-to-noise ratios than subordinate males (*dominant vs subordinate speed threshold event ratio*, Z = 3.98, *p* < .001; Figure 3d).

Finally, to summarize the differences between dominant and subordinate males in routine social contexts, we used principal component analysis (PCA) to combine trajectory and network metrics, finding that dominant and subordinate individuals clearly separate along PC1 (58.4% explained variance; Figure 3e).

## Discussion

In many species, cultures, and contexts, social dominance reflects the ability to control the behavior of group partners (16). Yet social influence is not unidimensional; it can be achieved in numerous ways, and the behavioral attributes that typify influence in one context may be the same that reduce influence in another. The question of what makes an effective leader is therefore not straightforward, and has different answers depending on the context in which leadership and influence are manifested.

Here we demonstrate that dominant male cichlid fish display behavioral traits frequently associated with leadership and increased social influence in many animal species including humans; such as higher aggression, social network centrality, and strong influence over routine group movement (12). Yet groups that contained dominant males as sources of information were slower to reach consensus in an association task. Rather, subordinate males, who occupy peripheral social network positions and have little influence over group movements, are the most effective in generating group consensus. This result was surprising considering that in other species such as rhesus macaques, subordinate males underperform a learned association task in the presence of more dominant males (29). Thus, the links between social *dominance* and social *influence* are not directly predictable, and highly context-specific.

The attributes that differentiated dominant and subordinate males, and likely their efficacy as agents of social influence, were social, spatial, and temporal. Dominant males occupied more central positions in interaction networks – a common trait of dominant individuals across species (9). This effect was driven by aggressive interactions of dominant males with other group members, whom they frequently chased and attacked. Group members continually fled from dominant males, and such aggressive behavior was rarely displayed by subordinate males. Consequently, dominant males had greater spatial separation from the rest of the group as they both chased away and were avoided by other group members, whereas subordinate males had higher spatial proximity to other group members (Figure 3b). This spatial separation likely led to lower opportunity for processes like physical leadership, spatial enhancement of stimuli, or observational learning. These differences may ultimately contribute to the slower time to group-consensus in groups with a dominant male informant. In contrast, the visual connectivity among naïve and informed individuals was not different for dominant and subordinate informants (Figure 3c). This suggests that visual access to social cues was similar for both subordinate and dominant males, potentially increasing the influence of dominant individuals (10). That dominant males had lower group influence, despite similar opportunity for visual attention, suggests the social cues from aggressive dominant males may have a negative valence compared to cues from passive subordinate males. In addition, other sensory modalities might contribute to dominant males’ potentially negative valence. For example, water-borne metabolites of circulating androgens, which are higher in dominant males, can signal dominance status to conspecifics (30). Similarly, mechanosensation via the lateral line has been shown to be another mode of non-contact social communication in *A. burtoni* (31).

Dominant individuals may also have been less reliable sources of social information due to their frequent, aggressive chasing of other group members. In the association task, informed individuals swam quickly and directly towards the ‘correct’ feeder, inducing following behavior in other group members. Such a rapid change in speed is also known to be an important social cue influencing collective behavior in zebrafish (32), and path speed and directedness known to affect both group coherence and information flow in golden shiners (33). However, this mode of swimming – rapid and straight – was a rare behavior for subordinate males, who typically moved at slow speeds without abrupt changes and stayed with the group. In contrast, dominant males frequently made rapid accelerations around the arena as they chased other individuals, which subsequently induced accelerated flight responses in subordinates. Thus, the frequent rapid movements of dominant males likely masked the informative social cues of a rapid swim towards the correct stimulus in the association task. This behavioral difference meant that dominant males had a lower signal-to-noise ratio of motion cues that served as sources of social information, likely contributing to their poor influence over group-consensus.

Overall, dominant males were behaviorally, spatially, and socially differentiable from subordinate males, consistent with many previous studies on this system (Figure 3e). These behavioral differences facilitated, and in fact defined, their dominant position, as well as their influence over normal group behavior. However, these same behaviors lowered their influence in a more sophisticated group task. This finding demonstrates that the attributes that define effective leaders may be highly context-specific, and traits commonly observed in socially dominant individuals may be the very traits that make them poor effectors of social change in other contexts. Thus, a passive process of leadership ascension, in which the most aggressive individuals rise to positions of influence, may be counter-productive in contexts where group consensus is prioritized.

## Materials and Methods

Captive *Astatotilapia burtoni* descended from a wild caught stock population (34) were maintained in stable communities until transfer to the experimental paradigm. Groups consisted of both males and females, though only males were used as informants since these have clear phenotypic indicators of social dominance, whereas females, although likely having social dominance hierarchies, have no reliable visual indicators of dominance status. All work was conducted in compliance with the Institutional Animal Care and Use Committee at The University of Texas at Austin.

### Behavioral paradigm

The behavioral task measured the time taken to reach group-consensus in a simple association task using a food reward and colored light-emitting diodes. Experiments were conducted in large, oval PVC tubs (205 liters, 108 × 54.6 × 42.7 cm) with two automatic fish feeders (EHEIM) mounted on opposite ends of each tank. The motor control pins of the feeders were rewired and externally controlled by a digital I/O switch slaved to an Arduino Uno microcontroller that also controlled one diffuse RGB LED mounted directly under each feeder (code available in supplementary material). Four times a day (0830, 1130, 1430, 1730) for five consecutive days, the Arduino randomly selected which fish feeder’s tumbler would turn, the LEDs simultaneously displayed one of two colors (RGB 255,60,0 [orange] or 0,255,255 [cyan]) for three seconds, followed by three seconds of no stimulus; then the tumbler of the feeder that displayed the orange stimulus would turn, spilling out a provision of Tetramin flake food. Neither of these color stimuli elicits an innate response in the focal animals (due for example to inherent color preferences;, 35), allowing their use as conditioned stimuli in an association learning paradigm. However, the color of the rewarded stimulus affected the speed at which the association was achieved, and we therefore kept the rewarded stimulus color consistent throughout all trials and randomized the location of colors to prevent spatial learning. A networked Logitech HD 1080p webcam was mounted above each tank and automatically scheduled to record for one minute before and after each training event using iSpy open source security camera software.

### Group-consensus task

Using the protocol described above, groups of eight *A. burtoni* (4 males, 4 females) underwent the training four times a day for five days. Group behavioral response to the task during all trials was scored as the proportion of individuals in the group that responded to the light stimulus by swimming towards it before the delivery of food. A successful group response was defined as seven or more of the eight group members swimming directly toward the positive stimulus in less than one second of stimulus onset, in two or more consecutive trials.

Within the five-day training period, all groups showed a behavioral shift from a lack of coordinated movement to a consensus movement toward the conditioned stimulus. After five days, all naïve groups reached consensus movement towards that correct cue, and subsequently one dominant male (“*dominant*”) and one subordinate male (“*subordinate*”) were placed into new groups (3 males, 4 females; total group size 8 individuals) that were naïve to the association task (Figure 1a). For groups with *dominant* males, all three other males were smaller than the *dominant*, while for groups with *subordinate* males, at least one male was larger than the *subordinate*. We did not observe any dominance shifts (i.e. a *dominant* becoming a *subordinate* in a new group, or vice-versa) in these group transitions. Seven groups each with either a *dominant or subordinate* informant were then placed in identical training protocols as previously and the time taken to group-consensus measured. In total, 168 fish were used.

### Deep-learning based automated tracking and analysis of behavior

We trained an implementation of a Mask and Region based Convolution Neural Network (Mask R-CNN) on a subset of manually labeled images to accurately detect and segment individual fish in the videos, resulting in pixel masks for each video frame and individual respectively (36–38). The masks were then skeletonized using morphological image transformations, allowing to estimate fish spine poses as seven equidistantly spaced points along the midline of each mask’s long axis. The first and second spine points represent head position and orientation, and were used to automatically reconstruct continuous fish trajectories using a simple, distance-based identity assignment approach. Accuracy and high detection frequency were visually verified with a Python-based GUI developed within the lab, that was also used to manually correct false identity assignments and losses (36).

### Behavioral, visual, and spatial connectivity analysis

In order to examine baseline differences in the behavior of *dominant* and *subordinate* males in social contexts, we placed seven additional groups of 10 individuals in identical tanks as described above and filmed their behavior in the absence of external stimuli (‘routine social context’). We calculated the behavioral, visual, and spatial interactions between all fish of each group. To estimate the number of behavioral interactions that dominant and subordinate males had with other group members, trajectory data was used to determine events with elevated swimming speed (above the 95% quantile of the speed distribution). The first two individuals passing this threshold in such events were treated as event initiator and responder, and a delay time between the two individuals was calculated (Figure 2c,d). We created behavioral interaction networks using the event count as the weights of the directed edges between network nodes (initiator to responder), that allowed the calculation of betweenness centrality (39)as a measure of behavioral influence in standard conditions (Figure 2). Additionally, the ratio of the sum of event durations to the recording’s duration was calculated for each fish, constituting the individuals’ hypothetical noise frequency in the social training context (fast, directed movement in absence of LED stimulus). Spatial connectivity between group members was calculated as their mean pairwise distances. Finally, we computed the visual connectivity as the mean angular area subtended by each individual on the retinas of all other group members, utilizing the contours of the Mask R-CNN detection results as occluding objects in a ray-casting approach (Figure 3a). Casting rays from both eyes of a focal fish towards to these contours (including the focal individual), we modelled the nearly complete field of view known from other freshwater fishes (40). These measures generated three connectivity scores for each dominant and subordinate group member: a behavioral (‘interaction’) connectivity, spatial (‘association’) connectivity, and visual connectivity. Finally, we conducted a principal component analysis (PCA) on the speed threshold event ratio (noise frequency) and connectivity scores to assess the overall consistency of metrics derived from trajectory and network analyses with the phenotypic indicators of dominance (Figure 3e).

### Data analysis and statistics

For time-to-consensus movement analysis, we conducted Kaplan-Meier Survival Analyses (41), using the first of two consecutive trials in which seven or more individuals responded to the stimulus onset as time to criterion, and right-censoring groups that did not complete the social learning task. We used log-rank tests to compare the survival estimates of naïve groups to informed groups, testing the effect of the informed individual’s social status. For comparisons of the baseline behaviors of dominant and subordinate fish, we performed multiple Wilcoxon signed-rank tests with the social status of group members as predictor and network centrality, mean angular area, mean pairwise distance, speed threshold event ratio and delay times as response variables. Here, a Bonferroni correction for n = 5 comparisons was applied.

## Acknowledgments

We thank members of the Jordan and Hofmann labs for many fruitful discussions. This work was supported by the NSF BEACON Center for the Study of Evolution in Action (AJ, HAH); Dr. Dan Bolnick and HHMI (AJ); the DFG Cluster of Excellence 2117 ‘Centre for the Advanced Study of Collective Behaviour’ (ID:422037984) (AJ, PN, OD); UT Austin Undergraduate Research Fellowships (LKG, SFG); a UT Austin Global Research Fellowship, UT Austin Graduate School Bruton and Summer Fellowships, and a Department of Integrative Biology Doctoral Dissertation Improvement Grant (MRS); and NSF grant IOS1354942 (HAH).

## Figures and Tables

**Supplementary Video 1.** A video showing the sequence of our tracking and behavioral analyses. The first sequence shows raw video input, upon a subset of which training annotations are performed. After training the neural network, fish in the video are then able to be automatically segmented with Mask R-CNN; this segmentation is here represented with a colored mask and corresponding bounding-box applied to each successfully detected fish in each frame of the video. Identities of each fish are maintained in subsequent frames using a nearest neighbor linking approach. The third visualization displays the trajectories and estimated spine positions of each individual. A simplified model (colored ‘fish’) is then overlaid on actual positions of fish to compute visual field connectivity using a ray-casting approach, considering the visual fields of each eye of a focal individual (gray-shaded areas) and occlusions (black-shaded areas).

